# Misleading inference of schistosome epidemiology from ribosomal internal transcribed spacer (ITS) and mitochondrial DNA

**DOI:** 10.64898/2026.04.30.721997

**Authors:** Egie E Enabuele, Roy N Platt, Ehizogie E Adeyemi, Martins S.O Aisien, Oluwaremilekun G Ajakaye, Mahmud U Ali, Ebube C Amaechi, Tolulope E Atalabi, Timothy Auta, Oluwaseun B Awosolu, Adamu G Dagona, Omoyemwen Edo-Taiwo, Chika P Ejikeugwu, Christopher Igbeneghu, Victor S Njom, Marian Onwude-Agbugui, Mary-Kate N Orji, Funso OP Oyinloye, Esther Oyemade, Habibat J Ozemoka, Christopher R Pam, Uchenna I Ugah, Jenna M Hulke, Grace A Arya, Timothy JC Anderson

## Abstract

The nuclear, internal transcribed spacer (ITS) and mitochondrial *cox1* markers are widely used to differentiate *Schistosoma haematobium* from its livestock counterparts, *S. bovis* and *S. curassoni*. *Schistosoma* isolated from humans with ITS and *cox*1 alleles from livestock parasites are typically inferred to be zoonotic infections and those with heterozygous ITS alleles (suggesting mixed species ancestry) are classified as recent hybrids. These classifications assume that the ITS and cox1 markers accurately reflect genome-wide ancestry. Here, we evaluated the reliability of this classification scheme by genotyping ITS and *cox1* from 132 parasites isolated from human urine, and from 37 adult schistosomes collected from cattle at 14 Nigerian locations. We also genome sequenced each sample to empirically determine livestock schistosome ancestry. ITS/*cox1* genotyping suggested extensive recent hybridization and zoonotic infection. Among parasites from humans, 10.1% carried both *S. curassoni* and *S. haematobium* ITS, consistent with F1 or early generation hybrids, 21% had livestock schistosome markers at both *cox1* and ITS suggesting zoonotic infection, while 13.7% carried *S. bovis cox1* alongside mixed *S. curassoni* and *S. haematobium* ITS, suggesting more complex ancestry. Genome sequencing revealed a very different picture. All parasites from humans formed a tight cluster regardless of ITS or *cox1* genotype, while all worms from cattle were well differentiated. We found no schistosomes containing 50% livestock parasite ancestry consistent with F1s. Instead, we observed regionally varying levels of *S. bovis* introgression, with modest levels in southern Nigeria (mean = 4.9%) and low levels in northern Nigeria (mean = 0.06%). These results demonstrate that: (i) two-locus genotyping is uninformative for detecting zoonotic infection or recent hybridization between *S. haematobium* and livestock schistosomes and (ii) previous data generated using this approach requires reinterpretation. These findings reveal the limitations of widely-used approaches for documenting zoonotic infection and hybridization between *S. haematobium* and livestock schistosome species.

**Author Summary:** Molecular markers provide useful tools for investigating epidemiology of closely related pathogens, but care is required when interpreting the data generated. This study critically examines the interpretation of two widely used molecular markers - the maternally-inherited mitochondrial *cox*1 locus and ribosomal internal transcribed spacer (ITS) – that are widely used for molecular epidemiology studies of schistosome parasites of humans and livestock. Data from these two markers have been used to infer recent hybridization between livestock and human parasites, and to identify livestock-to-human transmission. To examine the accuracy of epidemiological inferences from the *cox*1/ITS genotypes we collected parasites from both cattle and people in Nigeria. We genotyped each parasite using cox1 and ITS, as well as using whole-genome sequencing. *Cox*1/ITS genotyping suggested high rates of hybridization and livestock-to-human transmission. By contrast, whole-genome data show that parasites sampled from humans and cattle are distinct and correspond to independent species. The discrepant results demonstrate that genotyping just two markers provide insufficient resolution and can lead to erroneous conclusions about parasite epidemiology and poor management decisions. We suggest that future work should sample multiple markers from each parasite, using amplicon sequencing or other cost-effective approaches, to improve our understanding of schistosome epidemiology.

## Introduction

Interspecific crosses can be staged between many schistosome species in rodent hosts in the laboratory, and there is a rich literature describing this work [1, 2]. These studies have revealed limited barriers to interspecific mating, competitive mating asymmetries between schistosome species pairs, and potential for maintenance of some interspecific hybrids over multiple generations in rodent hosts. Furthermore, interspecific mating results in a spectrum of outcomes, ranging from production of fertile diploid progeny, to parthenogenic production of haploid progeny from the female parent [3] . There is also unambiguous evidence that interspecific hybridization occurs in nature between several pairs of species. For example, allozyme markers revealed evidence for hybridization between *S. haematobium* and *S. guineensis* during displacement of the latter species from much of Cameroon [4, 5], and revealed extensive hybridization between *S. haematobium* and *S. mattheei*, where these species are sympatric [6].

Much recent interest in schistosome hybridization has focused on interactions between the human schistosome *S. haematobium* and livestock schistosomes *S. bovis* or *S. curassoni* The first molecular evidence for *S. haematobium* with hybrid ancestry of *S. haematobium* came from field epidemiology studies 16 years ago [7], which revealed miracidia carrying *S. haematobium* ribosomal ITS and *S. bovis* mtDNA, as well as miracidia and cercariae with mixed *S. bovis*/*S. haematobium* profiles for ITS. There is now a burgeoning molecular epidemiology literature suggesting that hybridization occurs frequently [Supplemental Table 1; 3, 7, 8, 9-83]. These results have been interpreted as overwhelming evidence for frequent hybridization between *S. haematobium* and *S. bovis*, and this view has become widely accepted. High rates of hybridization between human and livestock schistosomes, if true, suggest that zoonotic spillover is frequent [46], alters our understanding of host–parasite epidemiology [47], and complicates disease management strategies [68]. Some have argued that *S. haematobium* transmission is best understood as a multi-host system [84, 85], similar to *S. japonicum*, in which multiple mammalian species, including water buffalo, rodents, and humans, serve as reservoirs [86].

Based on these findings, One Health control strategies for *S. haematobium*, including mass chemotherapy in both humans and livestock, are now being considered [84, 87] and are currently under discussion by the World Health Organization. Such approaches may influence how limited resources are allocated, so misclassification of levels of zoonotic infection and hybridization has significant programmatic implications. If zoonotic transmission is overestimated, control programs may divert resources toward livestock interventions that have little impact on human disease burden. Conversely, underestimating zoonotic transmission could leave animal reservoirs untreated, allowing continued human re-infection. Accurate identification of transmission sources is therefore essential for evidence-based policy.

How strong is the evidence for frequent zoonotic infection and natural hybridization between *S. haematobium* and *S. bovis* or *S. curassoni*? The most common method for identifying hybrids is to genotype two genetic markers in field-collected, parasite larvae hatched from eggs isolated from human urine samples [7], or from adult schistosomes collected from rodents [28] or livestock [81]. The two loci are the mitochondrial cytochrome oxidase 1 (*cox1*) gene and the nuclear internal transcribed spacer (ITS) rDNA region. The sequenced region of ITS spans parts of ITS1, 5.8S, and ITS2 of the 45S rDNA operon [Figure 1; 81]; we refer to this as ITS throughout the manuscript. Early comparative studies identified five positions within the ITS1 and ITS2 that were able to differentiate between a limited number of reference sequences. Heterozygous alleles at these 5 diagnostic sites or mito-nuclear discordance are often seen as evidence for natural hybridization [Figure 1; 81]. Similarly, identification of parasites infecting humans that carry livestock schistosome ITS genotypes and mtDNA genotypes, have been used to infer zoonotic infection [21, 44].

**Figure 1.**
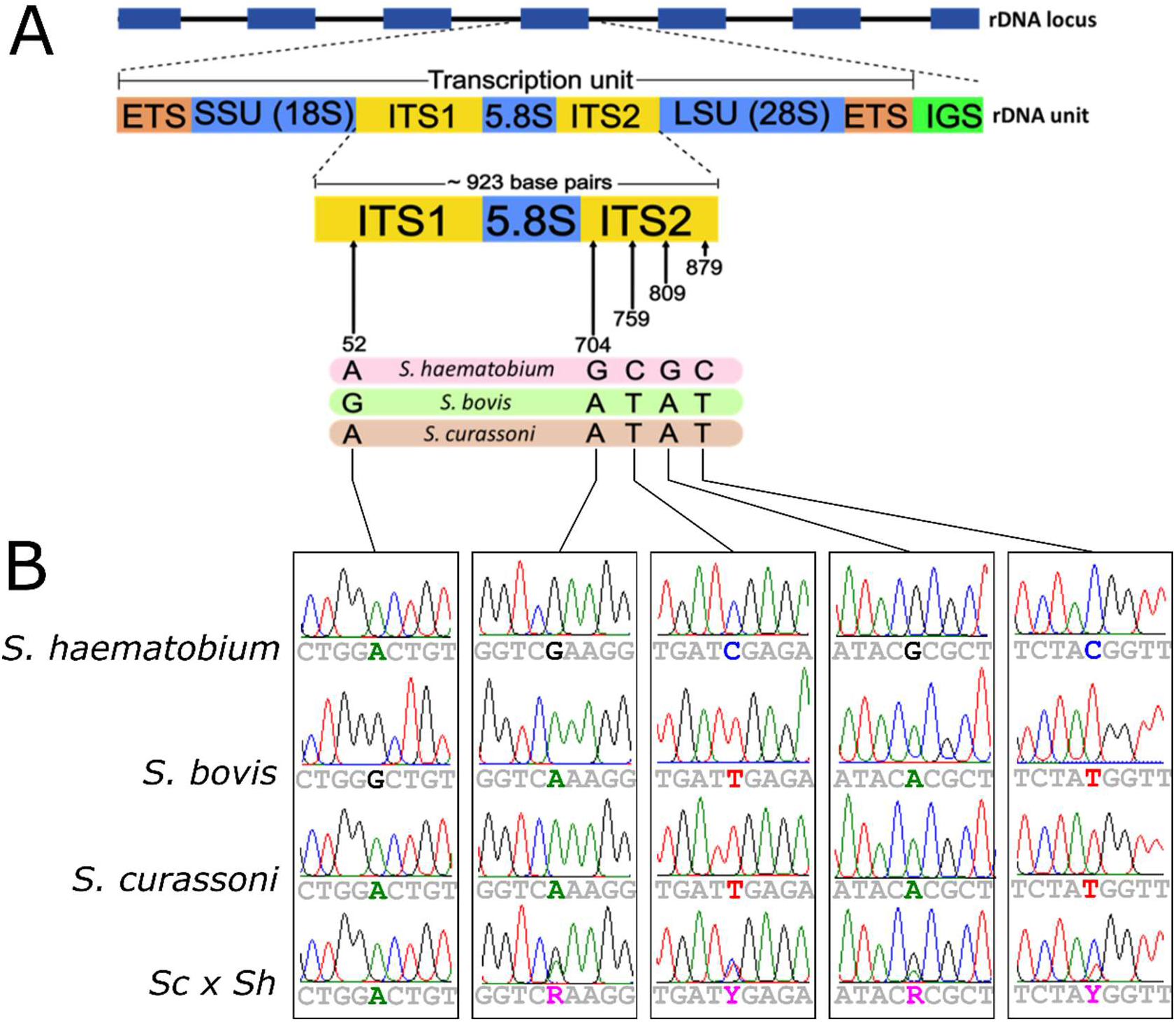
ITS structure and diagnostic alleles. (A) The internal transcribed spacer (ITS) is a region within the 45S rDNA operon that separates the 18S and 28S small and large ribosomal subunits. The ITS locus used for species identification in schistosomes spans the ITS1, 5.8S, and ITS2 regions. Five diagnostic sites within this locus are commonly used to classify schistosomes to the species level. (B) Sanger chromatograms at diagnostic positions along the ITS locus. Dual peaks at positions 704, 759, 809, and 879 indicate heterozygous alleles containing both *S. curassoni* and *S. haematobium* variants.

The criteria used to classify schistosomes as hybrids based on *cox*1 and ITS vary between studies. Schistosomes carrying discordant *cox*1 and ITS alleles – for example *cox*1 from *S. bovis* and ITS from *S. haematobium* - are often classified as “hybrids,” although this approach does not distinguish between recent hybridization events and those occurring hundreds of generations ago [62]. Other authors have used ITS genotypes to categorize hybrid *Schistosoma* [Table 1; 7, 26]. Parasites carrying ITS alleles from both *S. haematobium* and livestock *Schistosoma* species are assigned as F1 hybrids or early generation hybrids, while those with discordant ITS and mtDNA are classified as late generation hybrids [26, 46]. Importantly, these classification schemes have not been validated with genomic data so remain speculative.

**Table 1.**
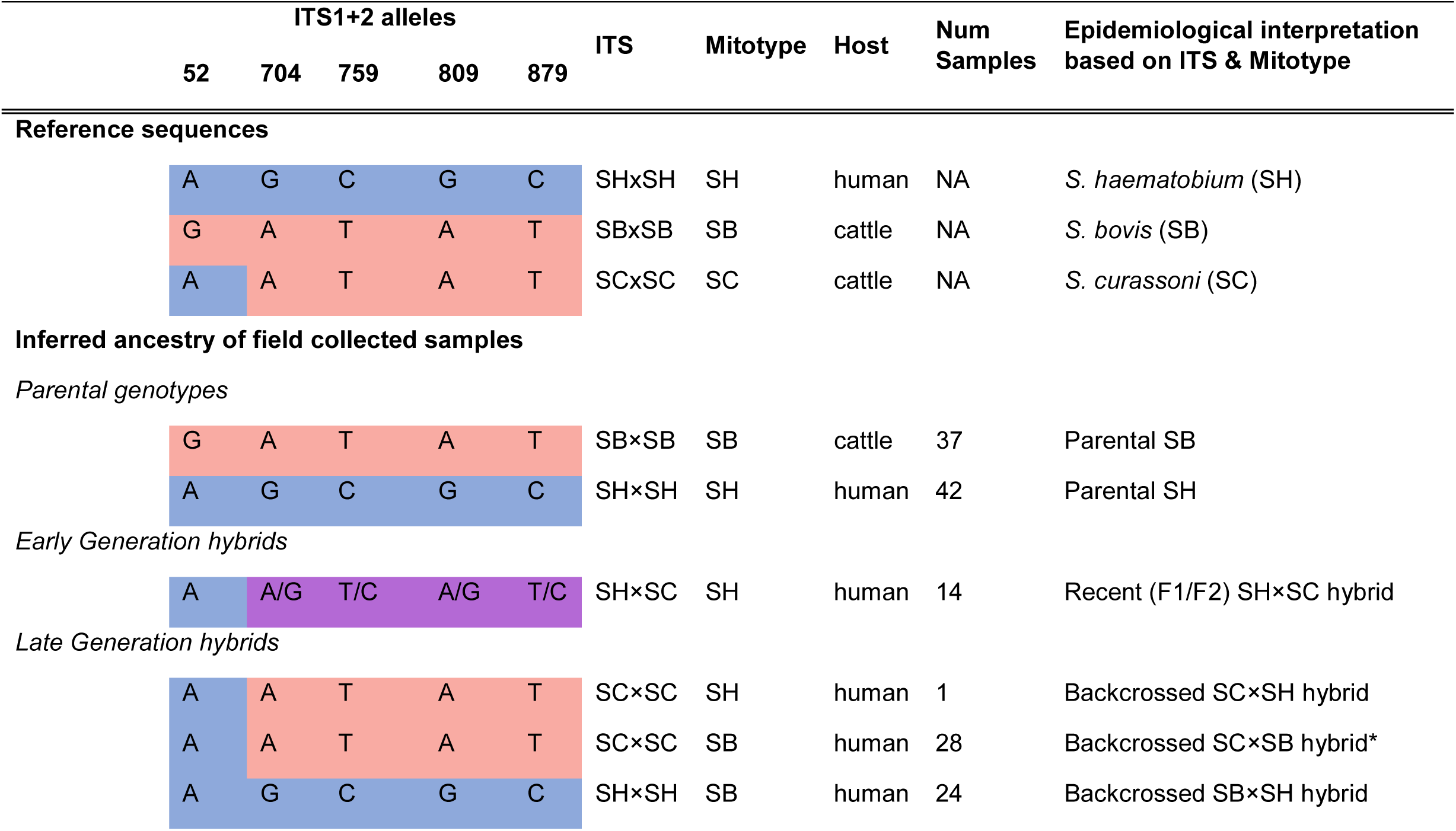
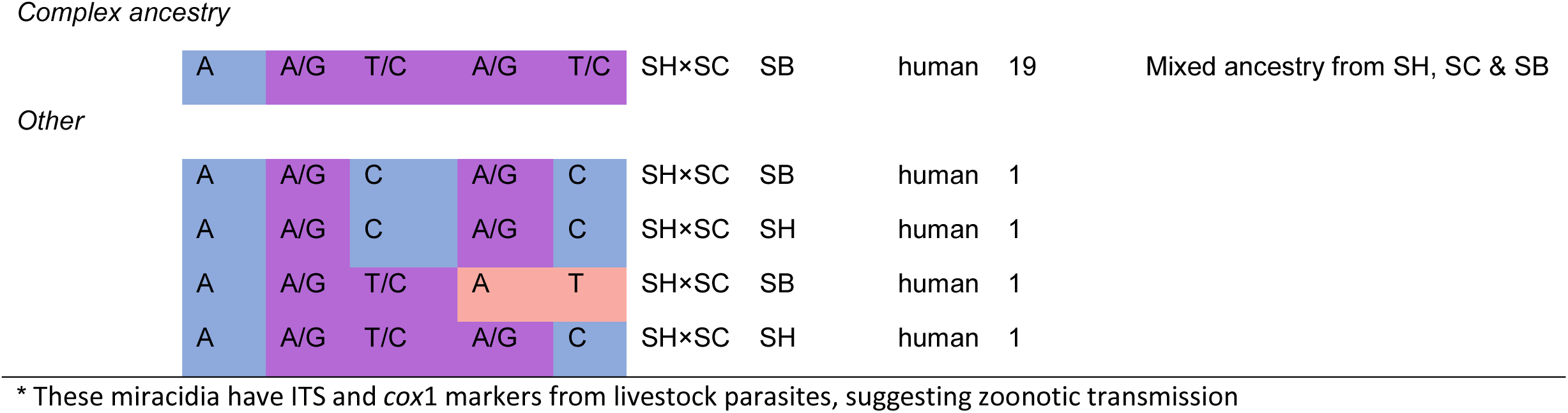
ITS and mitochondrial genotypes from Nigerian schistosome parasites. The ancestry of samples inferred from ITS and mtDNA marker scheme for *S. bovis* (“SB”), *S. curassoni* (“SC”) and *S. haematobium* (“SH”) following interpretations commonly used in the literature [46]. Only parasites collected from Nigeria are considered here. The interpretations from ITS/Cox1 genotyping are critically examined by comparison with whole genome sequence data in this paper but are not supported.

| ITS1+2 alleles |  |  |  |  | ITS | Mitotype | Host | Num Samples | Epidemiological interpretation based on ITS & Mitotype |
| --- | --- | --- | --- | --- | --- | --- | --- | --- | --- |
| 52 | 704 | 759 | 809 | 879 |  |  |  |  |  |
| Reference sequences |  |  |  |  |  |  |  |  |  |
| A | G | C | G | C | SHxSH | SH | human | NA | S. haematobium (SH) |
| G | A | T | A | T | SBxSB | SB | cattle | NA | S. bovis (SB) |
| A | A | T | A | T | SCxSC | SC | cattle | NA | S. curassoni (SC) |
| Inferred ancestry of field collected samples |  |  |  |  |  |  |  |  |  |
| Parental genotypes |  |  |  |  |  |  |  |  |  |
| G | A | T | A | T | SB×SB | SB | cattle | 37 | Parental SB |
| A | G | C | G | C | SH×SH | SH | human | 42 | Parental SH |
| Early Generation hybrids |  |  |  |  |  |  |  |  |  |
| A | A/G | T/C | A/G | T/C | SH×SC | SH | human | 14 | Recent (F1/F2) SH×SC hybrid |
| Late Generation hybrids |  |  |  |  |  |  |  |  |  |
| A | A | T | A | T | SC×SC | SH | human | 1 | Backcrossed SC×SH hybrid |
| A | A | T | A | T | SC×SC | SB | human | 28 | Backcrossed SC×SB hybrid* |
| A | G | C | G | C | SH×SH | SB | human | 24 | Backcrossed SB×SH hybrid |
| *Complex ancestry* |  |  |  |  |  |  |  |  |  |
| A | A/G | T/C | A/G | T/C | SH×SC | SB | human | 19 | Mixed ancestry from SH, SC & SB |

*Other*
|  |  |  |  |  |  |  |  |  |
| --- | --- | --- | --- | --- | --- | --- | --- | --- |
| A | A/G | C | A/G | C | SH×SC | SB | human | 1 |
| A | A/G | C | A/G | C | SH×SC | SH | human | 1 |
| A | A/G | T/C | A | T | SH×SC | SB | human | 1 |
| A | A/G | T/C | A/G | C | SH×SC | SH | human | 1 |

\* These miracidia have ITS and *cox1* markers from livestock parasites, suggesting zoonotic transmission

Here, we evaluate the evidence supporting recent hybridization and zoonotic infection by comparing inference from two-locus (*cox*1 and ITS) genotyping with empirical estimates of hybrid ancestry obtained through whole-genome sequencing. We genotyped 132 miracidia hatched from eggs isolated from human urine and 37 adult worms from cattle from 10 Nigerian states. We found that over half of the miracidia from humans exhibited mixed ancestry with either *S. bovis* or *S. curassoni* using *cox*1 and ITS. These included many with mixed ITS profiles that would be classified as recent hybrids, while multiple miracidia from humans carried livestock schistosome markers at both *cox*1 and ITS. By contrast, using 1.87 million whole-genome single nucleotide variants (SNVs), all miracidia from humans were well-differentiated from livestock schistosomes and provided no evidence for either recent hybridization or zoonotic infection. These results demonstrate that two locus genotyping using *cox1* and ITS is uninformative for species identification and for detection of recent hybrids and calls into question previous conclusions about recent hybridization between human and livestock schistosomes based on *cox*1 and ITS genotyping studies.

## Methods

### Ethics Statement

Ethical clearance for collection from humans was provided by National Health Research Ethics Committee of Nigeria (NHREC) (protocol number: NHREC/01/01/2007– 30/10/2020 and approval number: NHREC/01/01/2007– 29/03/2021) and by the IRB permit HSC-20180612H from the University of Texas Health Science Center. Informed written consent was obtained from all participants or their legal guardians.

### Sample collection

We collected 169 *Schistosoma* samples from Nigeria, including 132 miracidia hatched from eggs isolated from human urine. Schistosome eggs were collected from urine samples of infected children as previously described [88]. Mid-stream urine was collected in universal bottle containers between 10 am and 2 pm and transported to the laboratory for microscopic examination. We hatched eggs in fresh water by exposure to light and placed individual miracidia on Whatman FTA cards. This was done by an experienced parasitologist (EEE) using a two-step procedure: each miracidium was first transferred to a drop of fresh water and microscopically checked for presence of single miracidium before transfer to an FTA card.

We purchased fresh intestines of routinely slaughtered cattle from meat vendors at four abattoirs located in Akure, Auchi, Benin City, and Enugu in Nigeria. In the laboratory, the mesenteric vessels of the intestines were visually inspected for adult schistosomes. These were collected from the blood vessels surrounding the intestine using sharp tweezers, washed in saline solution, and stored in 80% ethanol.

### Single nucleotide variant (SNV) genotyping and analyses

We used whole-genome amplification (WGA) of single miracidia stored on FTA cards [89]. Adult schistosomes collected from cattle were sequenced following whole-genome amplification of 18-125 ng of DNA prepared from individual worms [89]. We prepared Illumina sequencing libraries following methods described in [89]. Briefly, DNA libraries were generated from 500 ng of WGA DNA per sample using the KAPA HyperPlus kit with the following modifications: (i) enzymatic fragmentation at 37°C for 10 minutes, (ii) adapter ligation at 20°C for one hour, and (iii) four cycles of library PCR amplification. Individual libraries were pooled at similar concentrations into a single library for sequencing. We sequenced the pooled library with 150 bp paired-end reads on an Illumina NovaSeq flow cell. We combined the newly generated reads with public data for combined downstream genotyping analyses. In addition, we included public sequence data from an additional 33 schistosomes. These were 9 *S. haematobium* [41, 63], 6 *S. bovis* [19], 7 *S. curassoni* [19], and 1 *S. margrebowiei* [90], as well as nine naturally occurring *S. bovis x S. curassoni* F1 and F2 hybrids, inferred from genome sequence data [19] and one laboratory-generated *S. haematobium × S. bovis* F1 hybrid [67]. Relevant information for all samples is provided in Supplemental Table 2.

We quality trimmed raw reads with Trimmomatic v0.39 [91] using the following parameters: LEADING:10, TRAILING:10, SLIDINGWINDOW:4:15, MINLEN:36, and ILLUMINACLIP:2:30:10:1:true. This removed low-quality bases at read ends, trimmed bases where quality fell below a Q15 threshold within a 4-nucleotide sliding window, and removed adapter sequences. Reads <36 nucleotides were discarded. We mapped trimmed reads to the Egyptian *S. haematobium* reference genome GCF_000699445.3 [92] using BBMap v38.18 [93] with the options ‘vslow’, ‘minid=0.8’, and ‘ambig=toss’. We assessed genome-wide mappability using GenMap v1.3.0 [94] with a 125 bp k-mer size and ≤ 1 mismatch. Mapped reads were sorted with SAMtools v1.13 [95], and PCR duplicates were marked using GATK v4.5.0.0 [96].

We called single nucleotide variants (SNVs) using GATK HaplotypeCaller and GenotypeGVCFs [96]. We hard filtered SNVs with VariantFiltration using: QualByDepth (QD < 2.0), RMSMappingQuality (MQ < 30.0), FisherStrand (FS > 60.0), StrandOddsRatio (SOR > 3.0), MQRankSum < –12.5, and ReadPosRankSum < –8.0. We removed sites failing minimum genotype quality (minGQ ≥ 30), site quality (minQ ≥ 30), or depth thresholds (minDP ≥ 10), multiallelic sites, indels, and variants on the ZW scaffold using VCFtools v0.1.16 [97]. We removed sites genotyped in <90% of individuals and individuals missing >20% of sites. These filters were chosen to ensure a reasonable number of makers were retained (>1 million) while maximizing the number of passable individuals. We exempted the laboratory-generated F1 *S. haematobium* (*Cameroon*) *× S. bovis* (Spain) hybrid [67] from the individual-level filtering despite its low read depth, because this sample provides the only positive control for a laboratory-generated first-generation *S. haematobium* X *S. bovis* hybrid.

### Mitochondrial analyses

We genotyped the mitochondrial genome using two approaches: (i) *<u>Cox</u>*<u>1 genotyping</u>: To replicate published studies, we used only the mitochondrial *cox*1 gene sequences for classifying mitotype. The *cox1* sequences for each sample were taken from *de novo* assembled mtDNA (see below)

(ii) *de novo* assembly of mtDNA: We used full length mtDNA sequences for each sample for more detailed phylogenetic analysis of mtDNA. We assembled mitochondrial genomes using methods described in [63] with minor modifications. A reference panel of mitochondrial genomes (Supplemental Table 3) was used to identify mtDNA derived reads, however *S. haematobium* (GCF_000699445.3) does not have a mitochondrial contig and previous work has shown that a large proportion of parasites in northern Africa, including Nigeria, contain an introgressed form of the *S. bovis* mitochondria [63]. We used the mitochondrial assembly from an *S. haematobium* collected in Angola to better represent a “true” non-introgressed mitochondrial haplotype. Assembled mitochondrial genomes were aligned with Clustal Omega v1.2.4 [98]. Reads were mapped to the reference mitochondrial panel with BBMap v38.18 [93] using ‘ambig=all’ and ‘minid=0.8’ to isolate mtDNA reads for *de novo*. Assemblies were generated using get_organelle_from_reads.py v1.7.7.1 [99] with 10 rounds of assembly and k-mer sizes of 21, 45, 65, and 85. Animal mitochondrial references (‘-F animal_mt’) guided the initial assembly rounds. Contigs were then scaffolded against the mitochondrial reference panel using RagTag v2.1.0 [100]. We selected and aligned the largest contiguous scaffold (>5,000 bp) from each sample with Clustal Omega v1.2.0 [98]. Low-confidence alignment regions were trimmed using trimAl v1.5.0 [101] and it’s ‘-automated1’ model. A mitochondrial phylogeny was inferred with RAxML-NG v1.2.2 [102] under the GTR substitution model, starting from 50 parsimony and 50 random trees, and using 1,000 bootstrap replicates. Branches with <50% support were collapsed using nw_ed v1.6 [103].

### ITS genotyping

We were unable to genotype ITS directly from the short-read data due to the highly repetitive nature rDNA arrays. Instead, we genotyped the ITS loci for all schistosome miracidia from humans and adult schistosomes collected from cattle, using the commonly used targeted PCR and Sanger sequencing techniques. This is the standard genotyping approach used in the literature. We used primers ETTS1 (TGC TTA AGT TCA GCG GGT) and ETTS2 (TAA CAA GGT TTC CGT AGG TGA A) [104] to amplify the ITS1–5.8S–ITS2 region (922 bp) from the same DNA aliquots used for Illumina sequencing. PCR cycling conditions included an initial denaturation at 95 °C for 5 min, then 40 cycles of denaturation at 95 °C for 30 s, annealing at 58 °C for 30 s, and extension at 72 °C for 90 s, and a final extension at 72 °C for 7 min.

Amplicons were confirmed on a 1% agarose gel and purified with ExoSAP-IT™ (ThermoFisher Scientific). Two PCR aliquots per sample were each combined with 5 µM of a forward or reverse primer and sent to Eurofins Genomics for sequencing.

We assembled and manually edited sequences using BioEdit v7.7.1.0 [105]. We called heterozygous peaks using the Indigo algorithm available on the Tracy web platform [106]. Sequences were trimmed by 50 bp from both ends to remove low-quality regions at the ends of each sequence. Heterozygous peaks were called when the secondary chromatogram peak heights reached >33% of the primary peak’s height. High-quality forward and reverse *ab1* files with unambiguous base calls were designated as reference profiles for *S. bovis* (GenBank Accession: PP546795), *S. haematobium* (GenBank Accession: PP546579), and *S. curassoni* (GenBank Accession: PP546674) for peak calling in Tracy [106].

### Two-marker classification using Mitochondria and ITS

We classified schistosomes into six categories based on mitochondrial cox1 and ITS genotypes following schemes adapted from Léger *et al* [Table 1; 46]. Three categories represented each “pure” parental species for schistosomes with homozygous ITS genotypes from *S. haematobium*, *S. bovis*, and *S. curassoni* and a concordant mitochondrial haplotype. “<u>Early generation Hybrids</u>” were heterozygous for ITS alleles regardless of mitochondrial type. For example, a schistosome heterozygous for *S. curassoni* and *S. haematobium* ITS alleles were classified as an early-generation hybrid. “<u>Late generation hybrids</u>” had homozygous ITS alleles from one species but discordant mitochondria from another (e.g., homozygous *S. haematobium* ITS with *S. bovis* mitochondria). “Complex” genotypes contained alleles from all three species, with heterozygous ITS sequences from two species and mitochondria from the third species. Samples not fitting any category were classified as “other”: these carried both homozygous or heterozygous SNPs at the 5 SNPs so haplotypes could not be determined. We then compared the inferred hybrid categories from two locus genotyping with empirical measures of hybrid ancestry determined from whole genome sequence data.

### Population genomic analyses

We examined genetic relationships among genome sequences using PCA, ancestry estimates, and phylogenetics. For PCA, linked SNVs with pairwise r² > 0.2 in 25 kb windows with a 5 kb step were identified using PLINK v1.90b6.2 [107] and removed with VCFtools v0.1.16 [97]. PCA was then performed using PLINK. For ancestry estimation, we used Admixture v1.3.0 [108] on the same PCA dataset but further thinned variants by removing any within 10 kb of each other. We used a supervised Admixture v1.3.0 [108] run with 1,000 cross-validation replicates to assess whole genome ancestry proportions based on a reference panel of three S*. haematobium*, three *S. bovis*, and three *S. curassoni* samples from outside Nigeria. Samples in the reference panel had concordant nuclear and mitochondrial genotypes, to ensure they represent ’pure’ parental ancestry when using a two-markers system. We estimated phylogenetic relationships using SVDquartets v1.49 [109] in PAUP* v4abuild168. To improve efficiency, we removed parsimony-uninformative SNVs with VCFtools. We analyzed 500,000 random quartets and used 100 bootstrap replicates to quantify nodal support. Branches with <50% bootstrap support were collapsed using nw_ed v1.6 [103].

### Statistical analyses

We examined variation in *S. haematobium* ancestry relative to sampling location, ITS genotype, and mitotype. *S. haematobium* ancestry was quantified as the Admixture component highest in non-Nigerian *S. haematobium* reference samples. We used an ordinary least squares model to assess the relationship between predictors and *S. haematobium* ancestry and to quantify their relative contributions. Model fit was evaluated using R² and F-tests relative to a null model.

### Computing environment and reproducibility

We performed all computational analyses on a standard desktop computer with ≥4 processors and ≥16 GB RAM or on a single computing node with 96 cores and 1 TB RAM. Command-line environments were managed using Conda v24.11.3. Environmental YAML files, Jupyter notebooks, and other code are archived at https://github.com/nealplatt/sch_hae_its. Alternatively, the files are permanently archived on Zenodo at https://doi.org/10.5281/zenodo.17807591.

## Results

### Study design

The central goal of this project was to compare results from two locus genotyping with ITS and mitochondrial *cox*1, with that obtained from whole genome sequencing. We genotyped ITS by PCR amplification and Sanger sequencing. This was done to replicate methods used in typical genotyping studies, and because directly genotyping ITS from Illumina sequence data is difficult. rDNA arrays are highly repetitive and mapping short reads to these regions usually fails. We found that the mappability of this region, measured with GenMap v1.3.0 [94], falls within the lowest 23rd percentile relative to the rest of the genome. We obtained both mtDNA and genome wide SNVs from Illumina sequencing data.

### Parasite collections and sequencing

We collected and sequenced 132 miracidia hatched from eggs isolated from human urine samples and 37 adult schistosomes from cattle from Nigeria for this study. We sampled miracidia from 53 patients (1-9 miracidia from each infected person) in 9 states and adult worms from 12 cattle from 4 slaughter houses in 4 different states. We generated high read depth (5.2-43x; median 43.7x) sequence from each sample. We included 34 sequences from previously published work [19, 41, 63, 90]. These included nine *S. haematobium*, six *S. bovis*, and seven *S. curassoni*, as well as nine natural *S. bovis × S. curassoni* hybrids [19], a laboratory-generated *S. haematobium × S. bovis* F1 hybrid [41], and a single *S. margrebowiei* sample used to root the analyses [90]. A description of each sample, along with collection details, accession numbers, and other metadata, is provided in Supplemental Table 2.

### ITS Sanger Genotyping

We genotyped ITS loci from bidirectional Sanger sequencing reads in 169 samples (132 miracidia from humans, 37 adult worms from cattle). All sequences are in GenBank (Supplemental Table 2) and Sanger chromatograms are available via Zenodo (https://doi.org/10.5281/zenodo.17807423).

We compared ITS genotypes at five commonly used diagnostic sites that distinguish *S. haematobium*, *S. bovis*, and *S. curassoni* alleles (Figure 1, Table 1). All parasites from cattle carried ITS with characteristic *S. bovis* SNPs. All parasites from humans carried either *S. haematobium (66/132; 50%)*, *S. curassoni* (29/132; 22%) haplotypes, or a mixed profile of both *S. haematobium* and *S. curassoni* haplotypes (37/132; 25%), while haplotypes could not be determined for 4 parasites (3%).

We identified 16 additional rare variable sites in ITS, which were conserved in ≥90% of samples. These 16 sites, together with the 5 diagnostic sites, defined 34 unique haplotypes (27 from human and 7 from cattle parasites; Supplemental Table 4). The most common genotype in parasites from humans was found in 44 ITS sequences from three countries (Madagascar, Nigeria, and Zanzibar). The most common haplotype among cattle parasites was found in 34 of 41 sequences from Nigeria, Côte d’Ivoire, and Uganda.

### Mitochondrial DNA Genotyping

We generated mitochondrial assemblies ranging from 12,756– 19,856 bp in length (mean = 14,109.8 bp). Mitochondrial genome lengths differed across species due to variants in long noncoding tandem repeats [110], so we used two rounds of alignment and trimming to ensure only homologous regions were compared for the mitochondrial phylogenetic tree (Figure 2). We extracted the *cox* 1 sequences from the assembled mitochondrial genomes to score mitotype. The *cox*1 alignment was 1,024 bp long and 73 haplotypes across all samples. Only 235 (23%) of sites were variable (Figure 2). Both the cox1 and mitochondrial genome trees recovered the canonical *S. haematobium* and *S. bovis* mitotypes in two well-supported clades, but the *S. bovis* clade contained two subclades of miracidia collected from humans. The presence of two distinct subclades suggests that the *S. bovis* mitochondrion has introgressed into *S. haematobium* populations on at least two independent occasions, rather than through a single historical event [63]. The raw *de novo* mitochondrial assemblies, trimmed alignment, and phylogeny are archived on Zenodo (https://doi.org/10.5281/zenodo.17807423).

**Figure 2.**
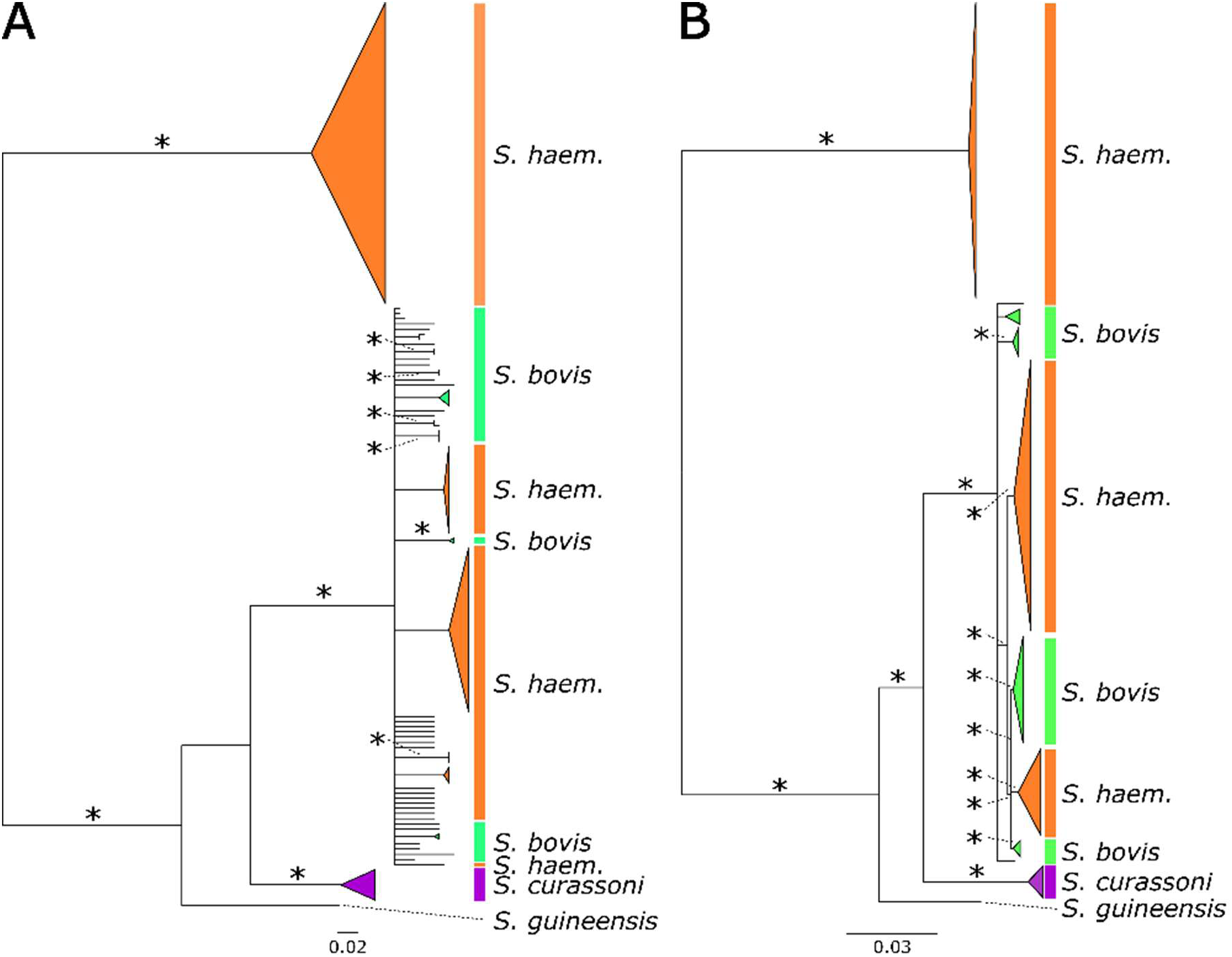
***cox*1 and, mitochondrial genome phylogenies.** The phylogenetic relationship between samples is affected by the marker type and number of available markers. The (A) *cox*1 and (B) mitochondrial genomes recover similar topologies with two major clades; a monophyletic *S. haematobium* clade and a polyphyletic group of *S. haematobium* and *S. bovis* samples. Resolution in the tree is increased as the number of markers increases from 1,024 in *cox*1 to 14,084 in the full mitochondrial genomes. Bootstrap values are shown for selected nodes, and nodes recovered in fewer than 50% of bootstrap replicates were collapsed.

### Categorization of *Schistosoma* samples using ITS/cox1

We used ITS genotypes and mitotypes to assign individuals to early or late generation hybrid classes following published classification schemes based on prior studies [26, 46]. All 37 adult worms from Nigerian cattle carried the *S. bovis* mitotype and homozygous *S. bovis* ITS alleles. By contrast, only 42/132 (31.8%) miracidia from humans carried *S. haematobium* mitotype and homozygous *S. haematobium* ITS consistent with pure *S. haematobium*. In total, 90/132 (68.2%) of miracidia from humans had either discordant nuclear and mitochondrial markers, heterozygous ITS alleles, or both.

Fourteen (10.6%) miracidia were categorized as early-generation *S. haematobium × S. curassoni* hybrids based on heterozygous ITS alleles. Twenty-five (18.9%) miracidia were categorized as late-generation hybrids. These included 24 parasites with homozygous *S. haematobium ITS* and *S. bovis* cox1 and one sample with homozygous *S. curassoni* ITS and *S. haematobium* cox1. Nineteen (14.4%) miracidia showed complex ancestry from all three species, containing heterozygous ITS from *S. haematobium* and *S. curassoni* and *S. bovis cox1*. Twenty-eight miracidia (21.2%) collected from humans carried homozygous *S. curassoni* ITS alleles and *S. bovis* mitochondria so had no evidence of *S. haematobium* ancestry. A final category contained four samples that were heterozygous at some, but not all, ITS diagnostic sites. Despite this ambiguity, all four miracidia carried some heterozygous *S. haematobium* and *S. curassoni* ITS alleles and either *S. haematobium* or *S. bovis* mitochondria.

### Whole-genome analyses

All reads generated in this study are available through the NCBI Short Read Archive (BioProjects PRJNA635500 and PRJNA561522). We genotyped 37,736,279 nucleotide sites in 210 *Schistosoma* samples. After filtering, 1,877,149 variants were genotyped in 203 individuals. These comprised 132 miracidia from humans and 37 adult worms from cattle from Nigeria generated in this study, and 34 sequences from previously published work (Supplemental Table 2). Each variant was genotyped in ≥193 samples, and each sample had ≥1,581,759 genotyped sites, with the exception of the laboratory-generated F1 *S. haematobium* × *S. bovis* hybrid, which was genotyped at only 247,864 of the 1.87M sites. This sample was retained due to its value as a true F1 *S. haematobium* X *S. bovis* hybrid control [41]. Descriptions of each sample, including read counts, genome coverage, mapping rates, and numbers of genotyped sites, are provided in Supplemental Table 2.

### Principal component analysis (PCA)

we removed physically linked sites, resulting in 278,855 unlinked autosomal SNVs. In the PCA, samples formed three major clusters along the first two PCs, which accounted for 62.2% and 22.6% of the variance in the SNV dataset, (Figure 3). One cluster corresponds to miracidia collected from humans, while parasites collected from livestock formed two additional clusters (*S. curassoni* and *S. bovis)*. The laboratory-generated *S. haematobium × S. bovis* F1 hybrid fell between *S. bovis* and *S. haematobium* as expected, despite missing 86.8% of genotypes. Field-collected F1 and F2 hybrids between *S. curassoni* and *S. bovis* identified in a previous publication [19] formed an intermediate continuum between the two parental species.

**Figure 3.**
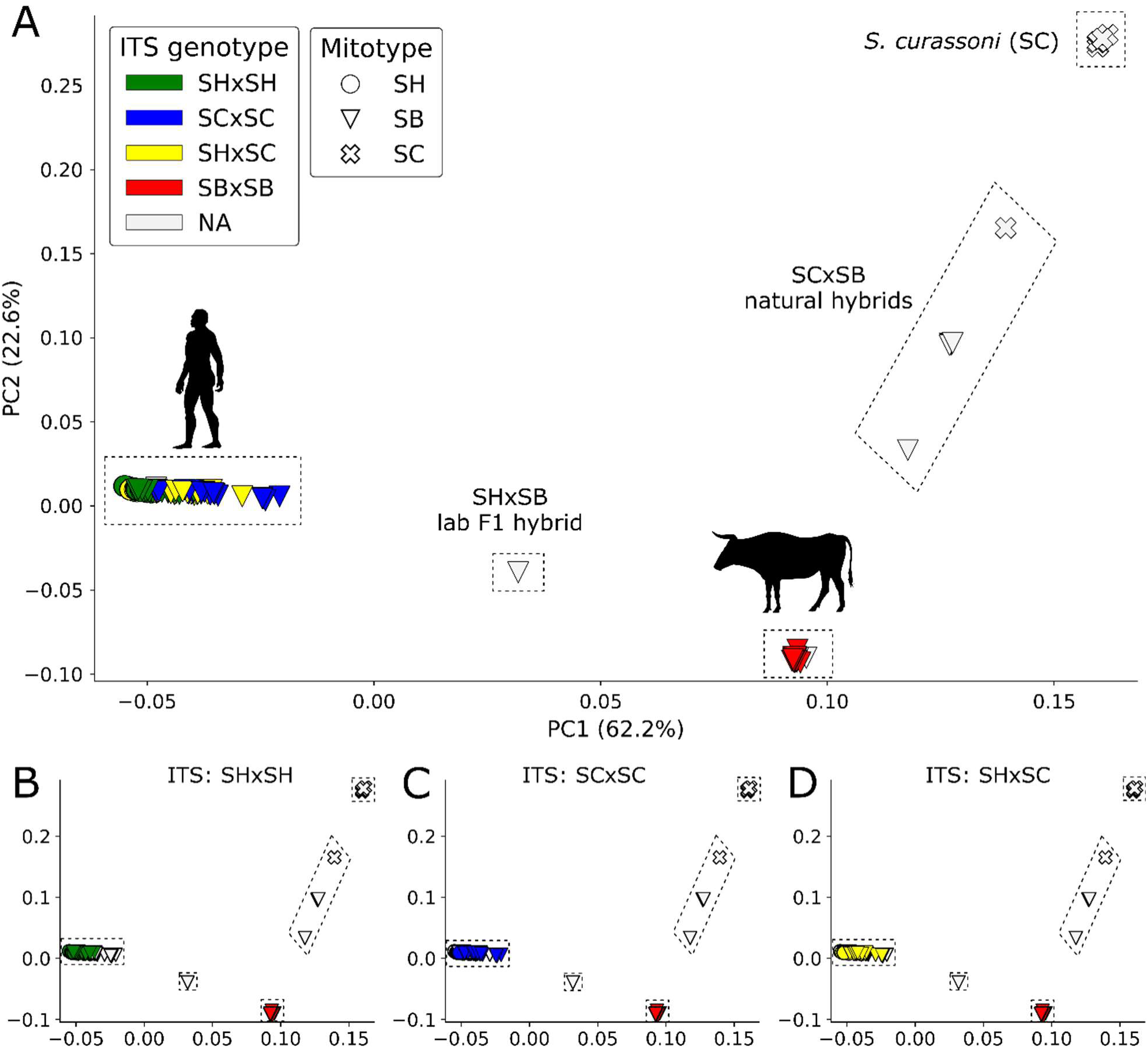
Principal component analysis (PCA) of 278,855 unlinked SNVs. The samples differentiated in genotypic space, as visualized with PCA. Each species forms a distinct cluster separated along PC1 and PC2. Known hybrid individuals fall intermediate to the parental species clusters, including the laboratory-generated F1 *S. haematobium × S. bovis* hybrid [67] and natural *S. curassoni × S. bovis* hybrids [19]. Samples are colored according to their ITS genotypes and marker shapes are used to identify the mitotype. All individuals collected from cattle presented homozygous *S. bovis* ITS alleles. Miracidia from humans presented *S. haematobium*, *S. curassoni*, or heterozygous *S. haematobium/S. curassoni* ITS alleles. None of the parasites with heterozygous *S. haematobium/S. curassoni* ITS alleles appear intermediate between *S. haematobium* and *S. curassoni*, as would be expected for early-generation hybrids. The three sub-plots show that all samples collected in humans form a single genetic cluster regardless of whether they carry (b) SHxSH, (c) SCxSC and (d) SHxSC, ITS alleles.

All adult worms collected from cattle in Nigeria fall in the *S. bovis* cluster. The miracidia collected from humans all fall in a single tight cluster. This is surprising given that the two-locus genotyping reveals miracidia from humans with ancestry from *S. haematobium*, *S. curassoni* and/or *S. bovis*. The two-locus genotyping identified 14 miracidia as recent hybrids between *S. curassoni* and *S. haematobium*. If true, we would expect these to fall intermediate between the *S. haematobium* and *S. curassoni* cluster, but they cluster with all other miracidia collected from humans. Similarly, the two-locus genotyping identified 28 miracidia with no *S. haematobium* ancestry: these carried *S. bovis cox1*, and homozygous *S. curassoni* ITS. We would expect these to cluster with livestock schistosomes in the PCA, but they cluster tightly with all other miracidia collected from humans. The two-locus genotyping also identified 19 miracidia with heterozygous ITS from *S. curassoni* and *S. haematobium, and S. bovis cox*1. We might expect these miracidia for which cox1/ITS indicate complex ancestry to be intermediate between *S. haematobium* and livestock schistosome clusters. However, the PCA clusters these parasites with all other miracidia from humans. Hence, there is strong disagreement between classification based on *cox*1/ITS, and PCA plots based on genome wide autosomal SNVs.

### Nuclear Phylogenetic analysis

We generated a coalescent-based phylogenetic tree using SVDquartets (Figure 4) and 1,061,546 phylogenetically informative autosomal SNVs. The resulting tree recovered monophyletic clades for each species, with only 12.74% of sampled quartets incompatible with the final topology. Nodal support across the tree was high (100%) at all nodes defining monophyletic species. ITS and *cox*1 genotypes are indicated at the branch tips. It is striking that miracidia from humans that are categorized as recent hybrids using ITS/cox1 genotyping, and miracidia that carry livestock schistosome markers at both ITS and cox1 cluster in a single monophyletic cluster containing all human derived miracidia.

**Figure 4.**
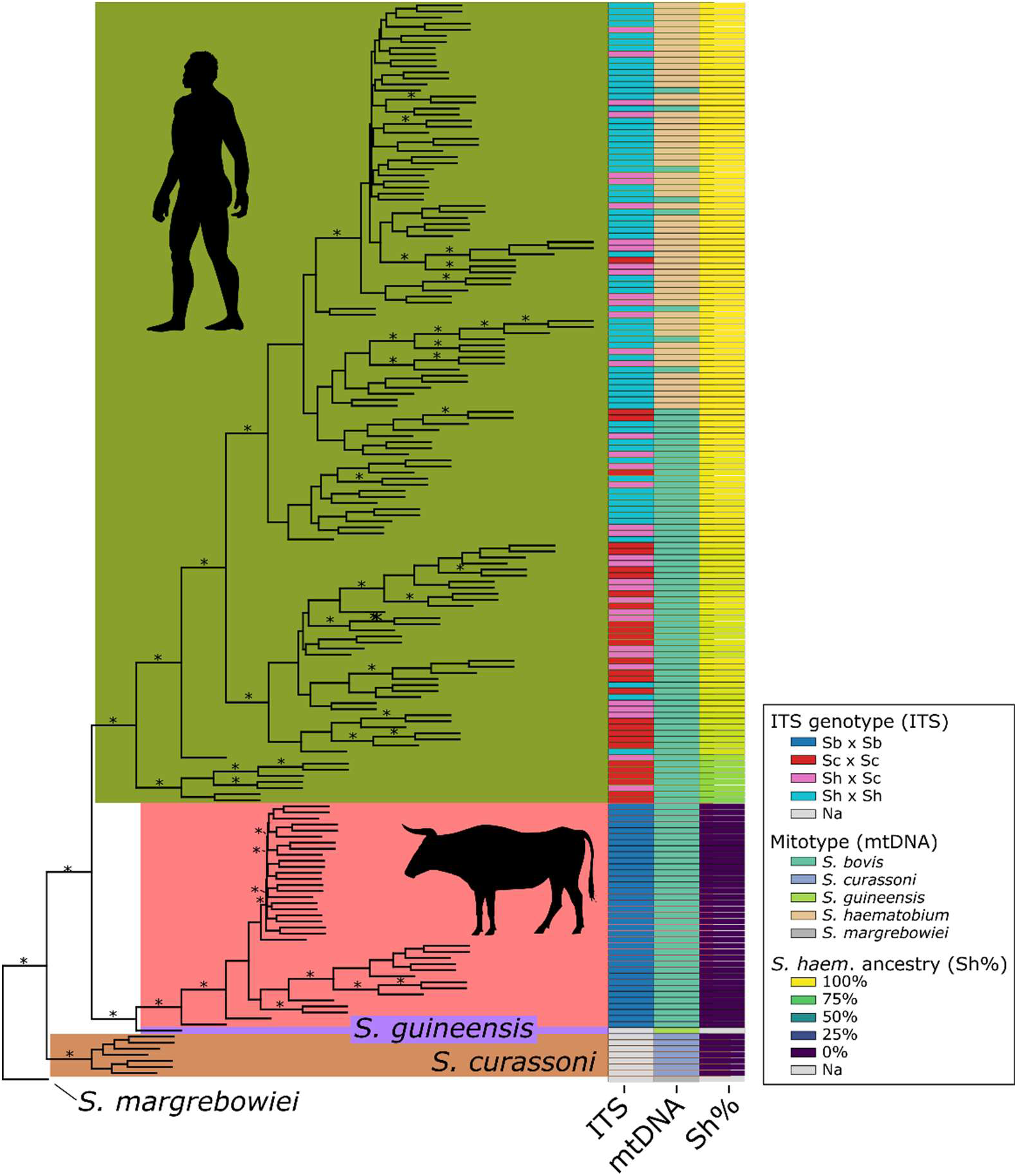
Whole-genome phylogeny from 1,061,546 phylogenetically informative, nuclear single nucleotide variants. A species tree generated with SVD quartets recovered monophyletic clades for miracidia sampled from humans and those sampled from cattle. Published *S. curassoni. S. guineensis* are included, and we used *S. margrebowiei* to root the tree. Mitotype, ITS genotype, and the *S. haematobium* ancestry component from the Admixture analysis are plotted at the tips of the tree. Mitotypes and ITS genotypes did not correspond consistently with clade structure. In contrast, the *S. haematobium* ancestry component clearly differentiates the *S. haematobium* clade from humans from the livestock *Schistosoma* clades. These results show that mitochondrial and ITS markers are not consistent with whole-genome ancestry patterns. Clades supported in ≥90% of bootstrap replicates are indicated with a “*”. Samples lacking an ITS are designated with “NA”. In each case these were samples sequenced in previous work. The human and cow silhouettes indicate the host from which each parasite was sampled.

### Admixture

We used nine individual samples as a reference panel for supervised Admixture analysis (Figure 5A) including three *S. bovis* (SRR11907383, SRR11907395, SRR11907458), three *S. curassoni* (ERR3012902, ERR5908628, ERR5919562) and three *S. haematobium* (SRR11907426, SRR11907431, SRR11907524). This analysis recovered distinct ancestry components within individual samples corresponding to *S. haematobium*, *S. bovis*, and *S. curassoni*, consistent with the PCA. The published, field collected, natural *S. bovis × S. curassoni* hybrids from cattle [19] contained 30–65% *S. bovis* ancestry consistent with F1 hybrids or F2 backcrosses. The laboratory-generated F1 *S. haematobium × S. bovis* hybrid contained 49% *S. bovis* and 51% *S. haematobium* ancestry, as expected, despite low genotyping success. These controls provide confidence in ancestry proportions estimated using Admixture.

**Figure 5.**
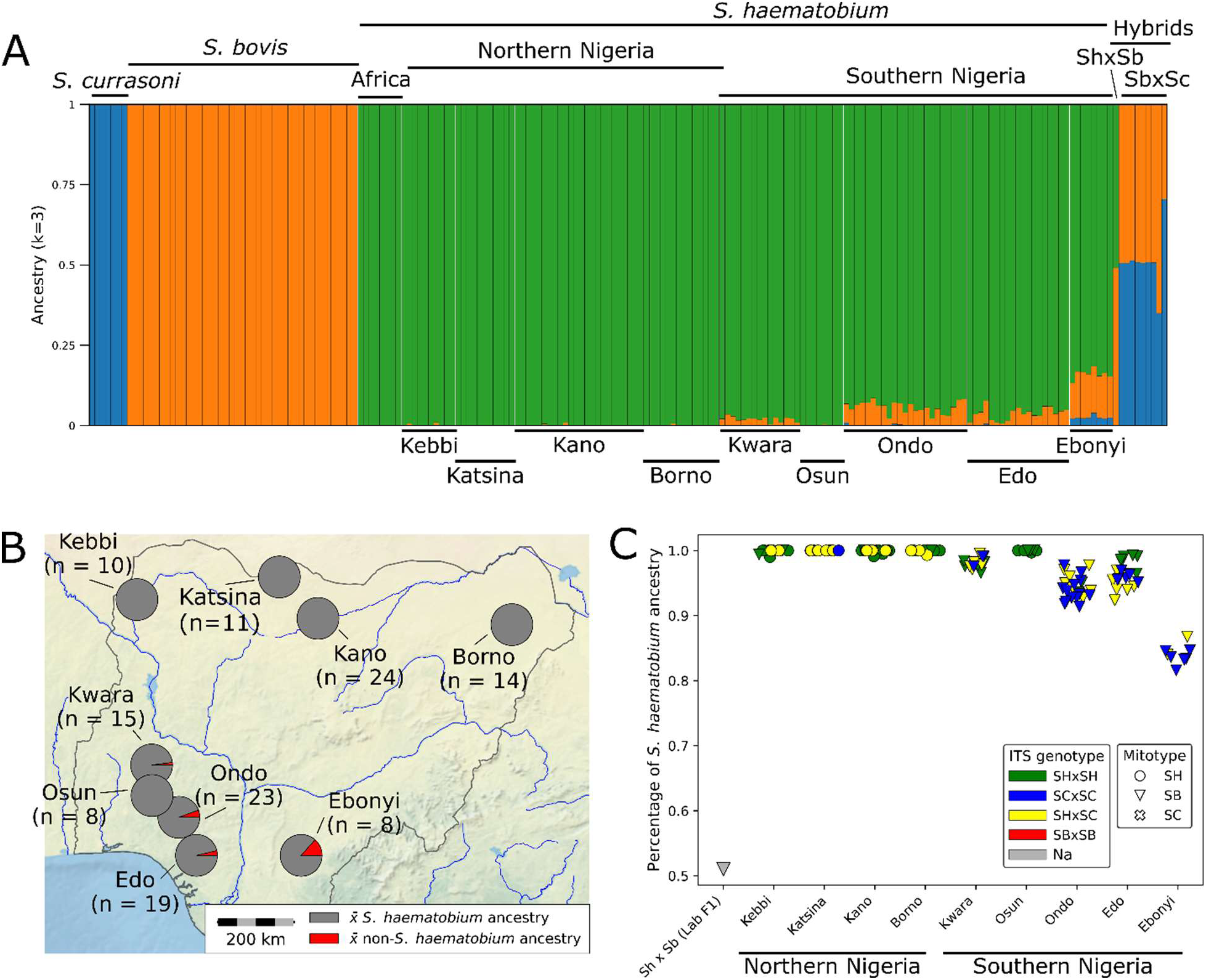
Biogeography of introgression across Nigeria. (A) A supervised Admixture analysis was used to estimate *S. haematobium* (green), *S. bovis* (orange), and *S. curassoni* (blue) ancestry in each parasite. Known hybrids—including an F1 *S. haematobium × S. bovis* hybrid [67] and both F1 and early backcross natural *S. curassoni × S. bovis* hybrids [19]—show high levels of mixed ancestry from their parental species. In contrast, we found no evidence of high levels of mixed ancestry in miracidia collected from humans. Instead, nearly all parasites collected in southern Nigeria carry low levels of livestock parasite ancestry, never exceeding 18.4% (mean 4.9%). “Africa” refers to published sequences from collection localities in Angola, Corsica, Cote d’ Ivoire, Madagascan, Namibia, Sao Tome, Senegal, Swaziland, Tanzania (Zanzibar), Uganda, and Zambia. (B) Mean *S. haematobium* ancestry values across sampling locations in Nigeria are shown on the map. Introgressed alleles are largely restricted to southern populations, which contain low levels of non–*S. haematobium* ancestry, with the exception of parasites from Osun. (C) The proportion of *S. haematobium* ancestry is relatively high across all Nigerian populations. This pattern is inconsistent with inferences from mitochondrial and ITS genotyping (Table 1). For example, individuals homozygous for *S. curassoni* ITS alleles and carrying *S. bovis* mtDNA still contain ∼97% *S. haematobium* ancestry across the nuclear genome.

Many of the Nigerian miracidia collected from humans carried low levels of *S. bovis* or *S. curassoni* ancestry. We refer to non–*S. haematobium* ancestry as “livestock ancestry,” since *S. bovis* and *S. curassoni* are both livestock parasites. On average, miracidia collected from humans harbored low levels of livestock ancestry (mean = range 0–18.5%), below expected ancestry proportions indicative of recent hybridization. In contrast, none of the adult worms from Nigerian cattle showed evidence of introgression.

None of the miracidia that are classified as recent hybrids using cox1/ITS contain 50% ancestry consistent with F1 hybrids. Similarly, miracidia from humans lacking *S. haematobium* markers at either ITS or cox1 are expected to show 100% livestock schistosome ancestry. However, these miracidia show low levels of livestock ancestry (mean = 7.17%, range: 0.002%-18.4%). The admixture analysis revealed no evidence for any recent hybridization between parasites from humans and livestock parasites, consistent with the PCA and phylogenetic analyses. We conclude that the genomic data provide a starkly different interpretation of schistosome hybridization and epidemiology when compared with cox1/ITS dataset.

### Biogeography of introgression in miracidia from humans

Levels of livestock introgression differed between locations (Figure 5B) but were remarkably similar in miracidia collected from people within a location (One-way ANOVA, F_8,123_ = 194.97, *p* = 4.6 x 10^-66^; Figure 5C). None of the *S. haematobium* parasites from northern states of Borno, Kebbi, Kano, Katsina or from the southern state of Osun, contained detectable introgressed alleles. In contrast, all but one parasite from Ebonyi, Edo, Kwara, and Ondo carried introgressed alleles at levels up to 18.4% (81.6% *S. haematobium* ancestry). Furthermore, all parasites samples from Ebonyi contained high levels of introgression (13.2%-18.4%, mean 16.0%), while parasites from Edo (0.7-7.6%, mean 3.7%), Kwara (0.5-3.5%, mean 2%) and Ondo (2.2-8.6%, mean 5.7%) contained less introgression.

Sampling sites can be categorized as northern versus southern. Introgression was restricted to populations in southern Nigeria, with the exception of Osun. Similarly, 68 of 73 parasites from southern populations carried introgressed *S. bovis* mitochondria. Osun was the only southern population with canonical *S. haematobium* mitochondria; in this location, 3 of 8 samples carried the introgressed *S. bovis* mitotype. In contrast, only 5 of 59 parasites from northern Nigeria carried the introgressed *S. bovis* mitotype. Finally, discordant mitochondrial and heterozygous ITS genotypes were more common in southern populations (75.4%) than in northern populations (28.8%) or in Osun (0%).

### Do ITS genotype and mitotype predict introgression?

Fig 5 reveals that *Schistosoma* samples that carry *S. haematobium* ITS, *S. curassoni* ITS or *S. haematobium*/*S. curassoni* ITS profiles in each location showed comparable levels of introgression in the nuclear genome. We used ordinary least squares regression to evaluate the contributions of geography, ITS genotype, and mitotype to *S. haematobium* nuclear ancestry (Table 2A). Geography alone was significantly associated with *S. haematobium* ancestry (model R² = 0.927, p = 4.6 × 10⁻⁶⁶). Adding ITS genotype and mitotype to a model that already included geography only marginally improved predictive power (ΔR² = 0.004; model R² = 0.931, p = 5.3 × 10⁻⁶⁴), and partial R² estimates indicated that ITS genotype partially contributes to model improvement (p = 0.048; partial R² = 0.051; Table 2B) and mitotype does not (p = 0.999; partial R² = <0.001; Table 2B). By comparison, geography alone accounted for ∼92% of the variance in *S. haematobium* ancestry. This result implies that individuals from the same location share similar levels of *S. haematobium* ancestry, regardless of mitochondrial and ITS genotypes.

**Table 2A.** Ordinary Least Squares model and ANOVA results.

| Model | Predictors | R<sup>2</sup> | $\Delta R^2$ vs.<br>Geography-<br>only | p-value |
| --- | --- | --- | --- | --- |
| 1 | Geography | 0.927 | — | $4.6 \times 10^{-66}$ |
| 2 | Geography + ITS | 0.931 | 0.004 | $4.1 \times 10^{-65}$ |
| 3 | Geography + mtDNA | 0.927 | <0.001 | $6.6 \times 10^{-65}$ |
| 4 | Geography + ITS + mtDNA | 0.931 | 0.004 | $5.3 \times 10^{-64}$ |

**Table 2B.** Ordinary Least Squares model & ANOVA results.

| Predictor | Partial R<sup>2</sup> | p-value |
| --- | --- | --- |
| Geography | 0.927 | $< 10^{-60}$ |
| ITS genotype | 0.050 | 0.0448 |
| mtDNA genotype | <0.001 | 0.999 |

## Discussion

### ITS and mtDNA do not identify F1 hybrids or zoonotic infections

This study was designed to evaluate the accuracy of *cox*1/ITS genotyping for identification of zoonotic infections and early generation hybrids between *S. haematobium* and livestock schistosomes by direct comparison with whole genome sequence data from the same samples.

Miracidia collected from humans carrying heterozygous *S. curassoni* /*S. haematobium* ITS and *S. haematobium cox*1 were common (14/132;10.6%) Nigeria. These miracidia would be inferred to be F1 or recent hybrids and therefore would be expected to carry 50% of their genome from both *S. haematobium* and *S. curassoni*. However, whole genome sequence data reveal that these putative F1 hybrid individuals showed low levels of introgression (mean = 0.04%, range: 0.002%-0.7%) from livestock schistosomes. Therefore, these miracidia result from old hybridization events and backcrossing, rather than F1 hybrids. Hence, the use of *cox*1 and ITS genotyping overestimates levels of recent hybridization in Nigeria.

Miracidia carrying heterozygous *S. curassoni* /*S. haematobium* ITS and *S. bovis* cox1 were abundant (19/132;14.4%). These miracidia carry markers from all three schistosome species, so might be expected to show complex ancestry. However, whole genome sequence reveals low levels of livestock schistosome introgression (mean = 5.2%, range: 0.5%-16%). Perhaps most surprising, miracidia with livestock markers at both ITS or *cox*1 were recovered from 28/132 (21.2%) miracidia from Nigerian urine samples from humans. These miracidia were homozygous for *S. curassoni* ITS and also carried *S. bovis cox1* and would therefore be expected to carry 100% of their nuclear genome from livestock parasites. However, whole genome sequence data unambiguously identifies these samples as *S. haematobium.* These putative zoonotic miracidia carried low levels of introgression from livestock parasites (mean = 7.17%, range: 0.002%-18.4%). Hence, genotyping of *cox*1/ITS incorrectly characterized these miracidia as progeny of zoonotic infections.

The unreliability of ITS/*cox*1 genotyping for identifying F1 hybrids, or zoonotic infections from livestock, lead us to investigate whether ITS/*cox*1 genotyping is informative for inferring levels of introgression in the schistosome nuclear genome. We found significant differences in levels of introgression between sampling locations in Nigeria. However, within each sampling location, parasites with different ITS or *cox*1 genotypes showed similar levels of introgression in the nuclear genome. We further formalized this analysis by using linear models to examine the predictive power of sampling location, *cox*1 and/or ITS genotype for predicting levels of *S. bovis* introgression. This revealed that sampling location was the strongest predictor of nuclear introgression. Adding ITS genotype to geographic location provided only marginal model improvements. *cox*1 genotype provided no power to predict levels of nuclear introgression. The simplest explanation for our results is that *S. curassoni* ITS alleles and *S. bovis* mitochondrial alleles are segregating within West African *S. haematobium* populations.

### Microsatellite studies from other West African countries

How broadly applicable are our results from Nigeria? Three other studies in Côte d’Ivoire, Senegal and Niger [16, 24, 60] have used genotyping of microsatellite loci to assess autosomal variation, in addition to *cox*1/ ITS genotyping. These three studies add strong support for our conclusions.

i. Angora *et al*. [16] examined 1966 miracidia collected from school children in Côte d’Ivoire using ITS2 restriction fragment length polymorphisms, *cox*1 genotyping and 10 microsatellite markers. Fifteen miracidia (0.7%) had an *S. bovis cox*1/*S. bovis* ITS2 genotype suggesting cross infection, while 407 miracidia had ITS2 profiles for both *S. haematobium* and *S. bovis*, consistent with F1 hybrids. However, they noted that the microsatellite markers revealed no differentiation between miracidia “presenting *S. haematobium*, *S. bovis* or hybrid profiles” for ITS2 and *cox*1 and concluded that “all genotyped parasites most likely belong to a single genetic entity”. We note that Angora et al genotyped ITS2 for which *S. bovis* and *S. curassoni* carry the same SNPs. It is therefore possible that these miracidia carried *S. curassoni* ITS rather than *S. bovis* as suggested. Miracidia carrying *S. curassoni* ITS are also commonly collected from humans in other west African countries including Mali [8] and Niger [44, 60].
ii. Boon *et al* [24] examined 730 miracidia collected from human urine samples from Senegal, and 62 adult *S. bovis* from cattle. They examined 186 of these using ITS, *cox*1 and 12 microsatellite markers. They identified 1 miracidium with ITS sequences from *S. bovis* and *S. haematobium,* suggesting an F1 hybrid, and 129 miracidia with *S. haematobium* ITS and *S. bovis* mtDNA. However, microsatellite markers clustered the putative F1 hybrid with the other parasites collected from humans, rather than intermediate between *S. haematobium* and *S. bovis* clusters. Boon *et al* [24] noted that miracidia collected from humans containing *S. bovis* mtDNA or ITS sequences “*appear to belong to a single randomly mating population*”.
iii. Pennance *et al* [60] genotyped single cercariae shed from snails in Niger using cox1, ITS and 6 microsatellite markers. They found cercariae with pure *S. bovis* profile (Sb *cox*1 and ITS), pure *S. haematobium* profile (Sh *cox*1 and ITS), as well as cercariae with mixed profiles (Sb Cox1 and Sh ITS, or Sb cox1 *S. haematobium*/*S. curassoni* ITS). However, PCA analysis using 6 microsatellites grouped all cercariae with pure *S. haematobium* or mixed profiles into one tight cluster that was well differentiated from the pure *S. bovis* cluster. The results from this study using just six microsatellites, closely mimic our results using genome wide SNPs, and suggest that parasites bearing heterozygous profiles at ITS are not recent hybrids.

The *cox*1/ITS two-marker system is currently used in most studies for identifying hybrid schistosome parasites (Supplemental Table 1). Results from such studies paint a picture of rampant hybridization and interbreeding between human and livestock parasites, directly contradicting results from the whole-genome analysis. Our findings, and those of Angora *et al*. [16] Boon *et al* [24], and Pennance et al [60] (see also [83]) demonstrate a need to re-evaluate putative F1 or early generation hybrid samples using additional genome wide markers.

### Implications for schistosome epidemiology and control

Our results suggest that *cox*1 and ITS genotyping overestimate levels of F1 hybrids and zoonotic schistosome infections. This is important because epidemiological conclusions from *cox*1/ITS genotyping studies have been used to argue for coordinated treatment programs of both humans and livestock to maximize efficacy of control, and to parameterize mathematical modelling studies of schistosome epidemiology [26].

Léger *et al.* [46] conducted one of the largest studies genotyping ITS and *cox*1 from 2,575 miracidia from 472 people and 115 ruminants from Senegal. Many of the miracidia from humans carried *S. bovis cox*1 together with *S. haematobium* ITS: these miracidia were assumed to result from old hybridization events, and multiple generations of backcrossing. However, they also identified 52 of 2575 (2%) miracidia with both *S. bovis* and *S. haematobium* ITS alleles, as well as *S. bovis* mtDNA, which they inferred to be *S. haematobium* - *S. bovis* F1 or early generation hybrids. Our results from Nigeria question this interpretation of their *cox*1/ITS data.

We speculate that most miracidia from Senegal [26, 46] with this profile will reflect historical introgression events rather than early hybrids. Genomic characterization of the miracidia inferred to reflect recent hybridization from the Léger *et al.* [46] and Borlase *et al.* [26] studies would allow accurate re-evaluation of the frequency of hybridization, and improve parameterization required for modelling of schistosome epidemiology as a multi-host system [26].

Two studies have reported evidence for suspected zoonotic infections of livestock parasites in humans: (i) Léger *et al*. [44] identified the same ITS/*cox*1 genotypes that we found in Nigeria with homozygous *S. curassoni* ITS and *S. bovis cox1*. They suggested that these genotypes resulted from backcrossing between *S. curassoni* and *S. bovis* in the human host. Similarly, Boissier *et al*. [21] examined *cox*1/ITS2 genotypes from 73 miracidia from 12 patients. They reported 1/73 miracidia with *S. bovis* ITS2 sequence and *S. bovis cox*1, and suggested that this finding “*presents the first potential evidence for the zoonotic transmission of S bovis*.” Genome-wide characterization of putative zoonotically derived miracidia from both Niger and Corsican study would be of particular interest. Our results from Nigeria suggest that these miracidia are unlikely to result from zoonotic infection. Furthermore, given that ITS2 sequences cannot distinguish *S. bovis* from *S. curassoni* (ITS1 one is needed for this), it is also possible that the Corsican parasites carry the *S. curassoni* ITS allele that is segregating at high frequency in Nigeria and in other West African *S. haematobium* populations [8, 44, 60].

### Population genetics of *S. haematobium* in Nigeria

Nigerian miracidia from humans show striking spatial variation in levels of nuclear introgression from livestock parasites. We observed that livestock-associated ancestry is largely restricted to miracidia collected from people in southern Nigeria, with the exception of Osun. Miracidia from people in Ebonyi, Edo, Kwara, and Ondo exhibit, on average, 5.5% non–*S. haematobium* ancestry. The highest levels of introgression occur in Ebonyi, where individual samples contain 13.2–18.4% non–*S. haematobium* ancestry. Additional genomic and molecular markers support a distinct biogeographic structure in southern Nigeria: introgressed *S. bovis* mitochondrial haplotypes are present in nearly every individual, and *S. curassoni* ITS alleles occur in three-quarters of parasites. The high frequency of *S. haematobium* carrying introgressed *S. bovis* mitochondria can explain the reports of human schistosomiases caused by *S. bovis* infections in Nigeria [11, 34].

In contrast, only 8.5% of individuals in northern populations carry introgressed *S. bovis* mitochondria, and 28.8% carry *S. curassoni* ITS alleles. Previous work in Nigeria identified two major *S. haematobium* genetic clusters, broadly corresponding to eastern and western populations [57]. All localities in Onyekwere *et al.* [57] lie south of Kaduna, Nigeria (10.5036° N, 7.4337° E), as do all sites in our southern population. Onyekwere *et al.* [57] reported high frequencies of introgressed *S. bovis* ITS alleles, whereas our results reveal *S. curassoni* ITS. The difference in conclusions are most likely because Onyekwere *et al.* [57] sequenced ITS2 only, while we sequenced both ITS1 and ITS2. *S. bovis* and *S. curassoni* ITS alleles differ by a single nucleotide difference in ITS1 (Fig 1). ITS2 contains four diagnostic positions that distinguish *S. haematobium* from both *S. bovis* and *S. curassoni*, but does not differentiate between the latter two species. Consequently, the ITS alleles detected in southern Nigerian populations in both this study and Onyekwere *et al.* [57] are likely to be *S. curassoni* reference alleles (see also <u>Implications for schistosome epidemiology and control)</u>.

### Consistent terminology and a clear definition of “hybrid” are needed to minimize confusion

Use of the vague blanket term “hybrids” to describe schistosome larvae carrying non-*haematobium* mtDNA or ribosomal DNA causes considerable confusion for three reasons: (i) Labeling individuals as “hybrids” may be misinterpreted by readers and policy makers as implying “early-generation hybrids,” even when authors do not intend this meaning. This gives the misleading impression that there is extensive mating and geneflow between *S. haematobium* and species infecting livestock; (ii) designating some miracidia as “hybrids” suggests that they are genomically distinct from non-hybrids. Yet, we show here presence that non-*haematobium* ITS or mtDNA are poor predictors of nuclear introgression levels. Furthermore, while *S. haematobium* sampled from across northern Africa may carry either *S. bovis* or *S. haematobium* mtDNA, almost all of these parasites have low levels of *S. bovis* introgression in the nuclear genome [63]; (iii) use of the nonspecific term “hybrids” may complicate comparison with other schistosomes species pairs, where interspecific crosses and F1 hybrids appear to be more common (e.g. *S. curassoni* x *S. bovis* [19]; *S. guineensis* x *S. haematobium* [111]). It is becoming clear that hybridization and introgression between schistosome species pairs presents a spectrum, ranging from rare to common. It is therefore important that suitable language used is available to reflect this. Webster and Huyse [83] make a parallel call for more nuanced terminology in this research field.

### Evolutionary perspectives

Previous work by our group and others now demonstrate that the *S. bovis* like mtDNA observed in many *S. haematobium* populations results from old hybridization events [62, 63, 67]. There are two lineages of *S. bovis* like mtDNA found in *S. haematobium* populations in samples from across northern Africa. The *S. bovis*-like mitochondria found in Nigeria fall into these two lineages. Hence, mtDNA does not provide a useful marker for recent hybridization. The current paper demonstrates that ITS genotyping is also a misleading marker for determining species identify and identifying early generation hybrids or zoonotic infections.

There are a number of biological explanations for the differences in ancestry observed at the ITS locus vs. the genome as a whole. These include lineage sorting and ancestral gene flow. Lineage sorting can occur in the absence of ongoing gene flow [112] when a polymorphism present in an ancestral population persists through multiple bifurcation (speciation) events. If the polymorphism is differentially fixed in the daughter lineages, the evolutionary history of the gene would not reflect true relationship among the daughter lineages and ancestral population.

Lineage sorting could result in *S. curassoni* ITS alleles segregating in *S. haematobium* populations without any hybridization. Incomplete lineage sorting is common. For example, 15% of genes in the human genome indicate that humans and gorillas are more closely related to each other than humans are to chimpanzees [113] as a consequence of lineage sorting. Across primates in general up to 64% of the genome is impacted by lineage sorting [114]. Within schistosomes, large-scale phylogenetics suggest a rapid emergence of multiple lineages in the *S. haematobium* group over a relatively short period of time [115]. This situation is likely to result in high-levels of lineage sorting [115]. If the five ITS alleles used as diagnostic markers for *S. bovis*, *S. curassoni*, and *S. haematobium* are not fixed in each species, lineage sorting could be misinterpreted as evidence of hybridization [112]. We note that the diagnostic ITS SNPs used for schistosome species identification were initially identified from a limited number of *S. bovis*, *S. curassoni* and *S. haematobium* specimens [81].

Regardless of the cause of the patterns observed, these results provide a cautionary tale about the dangers of inferring zoonotic infection or recent hybridization with insufficient marker loci [116]. Clinicians, epidemiologists and public health officials should be aware that the presence of livestock-associated markers in a patient sample does not necessarily indicate a zoonotic infection or a recent hybrid but may simply reflect ancient introgression that is common in some populations. Two markers are clearly insufficient for identifying hybridization or zoonotic transmission, and whole genome sequence is cost prohibitive. However, inexpensive methods for genotyping several hundred markers across the genome can provide a scalable alternative for future studies of schistosome hybridization and population genetics [64]. For example, amplicon panels allow genotyping of multiple loci and sequencing costs are minimized because hundreds of samples can be pooled together in a single run. Adoption of such methods will greatly reduce confusion in this field and improve our understanding of schistosome epidemiology and control.

### Study Limitations

The study is geographically limited to Nigeria; the extent to which these findings apply to other West African countries or to other parts of Africa remains to be determined. Whole-genome sequencing, while informative, was only performed on 132 miracidia from humans. Genome sequence data revealed no evidence for hybridization or zoonotic infection in these samples, in contrast to the ITS/cox1 genotyping. However, deeper sampling may potentially reveal true hybridization events or evidence for zoonotic transmission.

## Acknowledgements

We thank Sandy Smith and John Heaner for providing computational support through the Texas Biomedical Research Institute High-Performance Computing Cluster. Comments from Bonnie Webster, David Rollinson and Eric Loker improved the manuscript. This research was funded by the National Institute of Allergy and Infectious Diseases (NIAID R01 AI166049-01 to TA). The funders had no role in study design, data collection and analysis, decision to publish, or preparation of the manuscript

## Supporting Information Legend

*SData-1_its_ab1_chromatograms.zip* – Sanger chromatograms for all ITS sequences in the .abi format.

*SData-2_reference_mitochondria.fasta* – FastA file containing mitochondrial genomes from public data. All sequences are available through NCBI or as supplemental data in previous work. These sequences were used as species-level reference mitochondria seed mitochondrial assemblies.

*SData-3_mito_assemblies.raw.fasta* – FastA file containing *de novo* mitochondrial genome assemblies using GetOrganelle. Assemblies are in their raw state without any filters applied.

*SData-4_mito_assemblies.aligned.trimmed.fasta* - FastA file containing *de novo* mitochondrial genome assemblies that have been aligned with Clustal Omega v1.2.0 [98]. Low-confidence alignment regions were trimmed using trimAl v1.5.0 [101] and it’s ‘-automated1’ model.

*SData-5_mito_assemblies.aligned.trimmed.fasta.raxml.support.nwk* – A mitochondrial phylogeny inferred with RAxML-NG v1.2.2 [102] under the GTR substitution model, starting from 50 parsimony and 50 random trees. Nodal support is provided as percentages from using 1,000 bootstrap replicates

